# Long-Term Leisure-Time Physical Activity and Other Health Habits as Predictors of Objectively Monitored Late-Life Physical Activity – A 40-Year Twin Study

**DOI:** 10.1101/205856

**Authors:** Katja Waller, Henri Vähä-Ypyä, Timo Törmäkangas, Pekka Hautasaari, Noora Lindgren, Paula Iso-Markku, Kauko Heikkilä, Juha Rinne, Jaakko Kaprio, Harri Sievänen, Urho M. Kujala

## Abstract

**IMPORTANCE:** Moderate-to-vigorous physical activity (MVPA) in old age is an important indicator of good health and functional capacity enabling independent living.

**OBJECTIVE:** To investigate whether physical activity and other health habits at ages 31-48 years predict objectively measured MVPA decades later.

**DESIGN, SETTING, AND PARTICIPANTS:** This prospective twin cohort study in Finland comprised 616 individuals (197 complete twin pairs, including 91 monozygotic pairs, born 1940-1944), who responded to baseline questionnaires in 1975, 1981, and 1990, and participated in accelerometer monitoring at follow-up (mean age, 73 years).

**EXPOSURES:** Primary exposure was long-term leisure-time physical activity, 1975-1990 (LT-mMET index). Covariates were body mass index (BMI), work-related physical activity, smoking, heavy alcohol use and health status in 1990, and socioeconomic status.

**MAIN OUTCOMES AND MEASURES:** Physical activity was measured with a waist-worn triaxial accelerometer (at least 10 hours per day for at least 4 days) to obtain daily mean MVPA values.

**RESULTS:** High baseline LT-mMET index predicted higher amounts of MVPA (increase in R^2^ of 6.9% after age and sex adjustment, *P*<.001) at follow-up. After addition of BMI to the regression model, the R^2^ value of the whole multivariate model was 17.2%, and with further addition of baseline smoking, socioeconomic status, and health status, the R^2^ increased to 20.3%. In pairwise analyses, differences in MVPA amount were seen only among twin pairs who were discordant at baseline for smoking (n=40 pairs, median follow-up MVPA 25 *vs*. 35 min, *P*=.037) or for health status (n=69 pairs, 30 *vs*. 44 min, *P*=.014). For smoking, the difference in MVPA also was seen for monozygotic pairs, but for health status, it was seen only for dizygotic pairs. Mediation analysis showed that shared genetic factors explained 82% of the correlation between LT-mMET and MVPA.

**CONCLUSIONS AND RELEVANCE:** Low leisure-time physical activity at younger age, overweight, smoking, low socioeconomic status, and health problems predicted low MVPA in old age in individual-based analyses. However, based on the pairwise analyses and quantitative trait modeling, genetic factors and smoking seem to be important determinants of later-life MVPA.

## Introduction

Reduced physical activity in old age predisposes strongly to disability while exercise-based rehabilitation improves measured and self-rated function among individuals with various chronic diseases,^1^ and prevents disability at older ages.^2^ High participation in moderate-to-vigorous physical activity (MVPA) at older ages is an indicator of good physical fitness and health, and consequently predicts reduced risk of disability and death in the older population.^3,4^

Some observations suggest that midlife low physical activity, obesity, and poor health status predict sedentary behavior in old age.^5^ Low physical activity^6,7^ and other lifestyle factors, such as smoking and use of alcohol^8–10^ predict or are associated with later disability and impaired mobility. However, no data exist describing whether long-term leisure-time physical activity during adulthood predicts objectively measured physical activity/mobility in old age. Non-communicable diseases and performance and activity limitations develop slowly, so it is important to investigate the long-term predictors of later-life physical activity levels.

Twin, family, and molecular genetic studies provide evidence for a role of genetic factors in obesity, many non-communicable diseases, fitness, and participation in physical activity, but the identity of specific genes for physical activity remains largely unknown.^11^ Thus, both genetic factors, including the possibility of genetic pleiotropy, and childhood environment-related factors may predispose to different clusters of risk factors and associated diseases.^3,12^ By studying outcomes in twin pairs discordant for exposure to different health habits and health outcomes, the possible confounding role of genetic and shared early childhood experiences can be considered. Twin pairs almost always share the same childhood family environment. Dizygotic (DZ) pairs share, on average, half of their segregating genes (like non-twin siblings), while monozygotic (MZ) pairs are genetically identical at the sequence level. Co-twin control analyses among discordant MZ twin pairs allow for stronger estimates of causal influences compared to associations seen in unrelated individuals.

We investigated how self-reported long-term leisure-time physical activity and other health habits from ages 31 to 48 years predict objectively measured physical activity and sedentary behavior at a mean age of 73.

## Methods

This MOBILETWIN study is an ancillary to the older Finnish Twin Cohort Study.^13^ Written informed consent was obtained from all participants, and the study was approved by the Ethics Committee of the Hospital District of Southwest Finland on 20 May 2014.

### Participant Inclusion

The study is based on a nationwide sample of all same-sex twin pairs born before 1958 with both co-twins alive in 1975.^13^ A baseline questionnaire was sent to all twin candidates in 1975. Among those whose home addresses could be identified (93.5%) in 1975, the response rate for twins was 87.6 %. A subsequent questionnaire was mailed in 1981 to all of the verified twins. The corresponding response rate among those responding in 1975 and alive in 1981 was 90.7 %. A third questionnaire was sent out in 1990 to all twin individuals aged 33-60 (birth cohorts 1930-1957) years who had responded to at least one of the earlier questionnaires (response rate was 77.3% of all surviving cohort members).^14^

For the current physical activity study (MOBILETWIN), twins from the 1940-1944 birth cohorts were selected (Figure 1). Altogether, 3186 twin individuals belonged to these birth cohorts and had responded to at least one of the first two questionnaires (1975 or 1981). A total of 145 twin individuals were excluded because they had participated in one of the previous studies on psychiatric disorders (schizophrenia and bipolar studies). All remaining 816 complete twin pairs, i.e., both alive and contactable, were invited to participate in the present study for a total of; 256 MZ, 490 DZ and 70 with unknown zygosity. The twins were sent an invitation letter in which they chose whether to participate in a health and cognition telephone interview and/or accelerometer study complemented with physical functioning questionnaire. Altogether, 1012 (61.9%) twin individuals participated in the telephone interview, 791 twin individuals wore the accelerometer for the required time, and 817 individuals filled in the whole questionnaire on physical functioning. A total of 616 participants (197 complete pairs, including 91 MZ and 95 DZ pairs) in the accelerometer study also had baseline physical activity data for all the baseline time points (1975, 1981, and 1990). For other baseline health variables, we maximized the statistical power of the analyses by including all possible twin individuals and discordant twin pairs who had data for these other health habits; therefore, the number of participants in different analyses may have varied according to variable under investigation.

**Figure 1.**
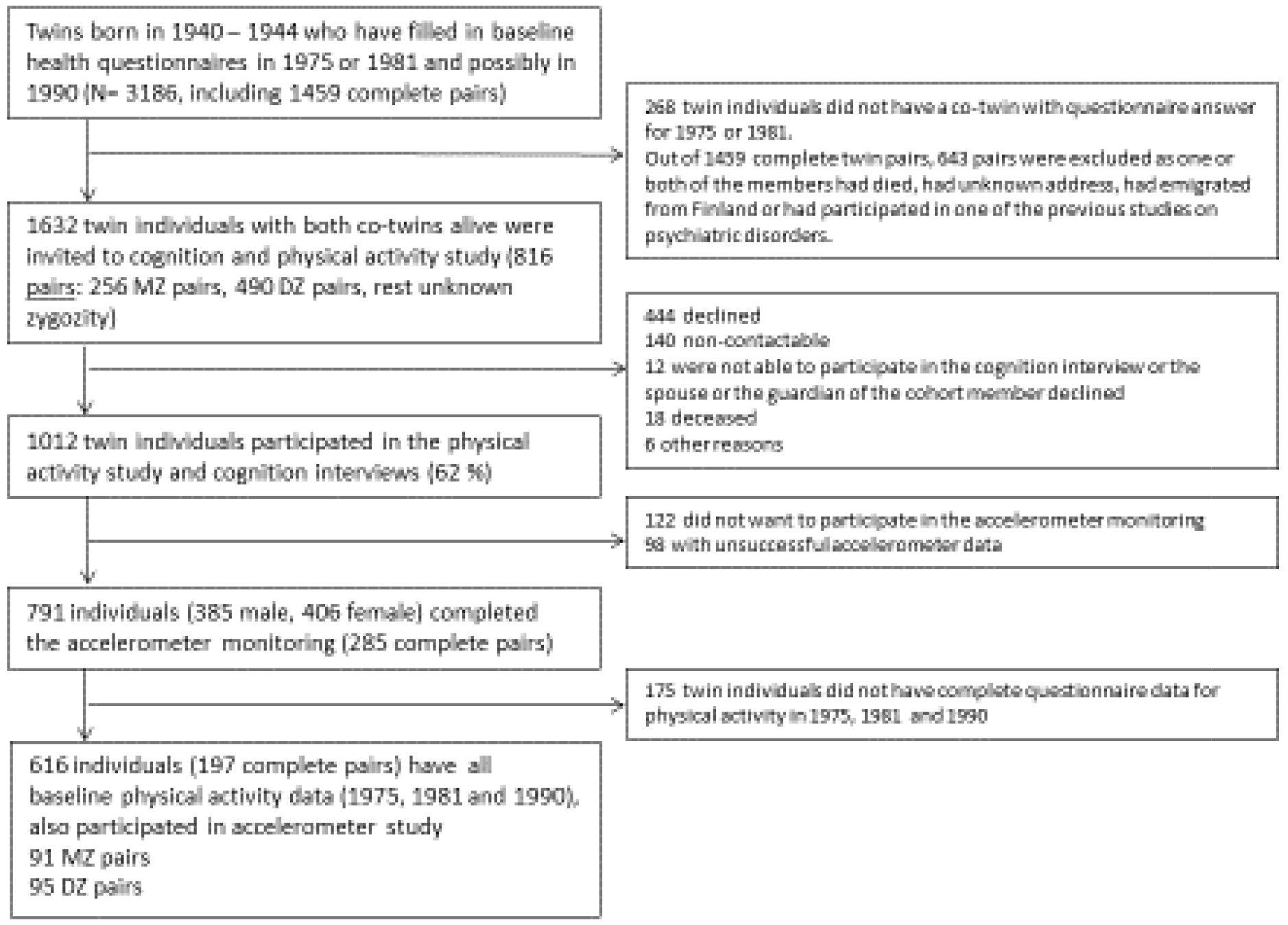
Participant Flow Diagram

### Baseline Predictor Assessment

The postal questionnaires in 1975 and 1981 were very similar, but the questionnaire in 1990 was slightly different in some parts; however, they all included questions on physical activity, occupation, work-related physical activity, smoking, use of alcohol, and physician-diagnosed diseases (available on the Twin Study website: http://www.twinstudy.helsinki.fi).

*Physical activity* habits were assessed by identical questions in 1975 and 1981 and with slightly different questions in 1990. All three questionnaires enabled calculation of the MET index. On the bases of earlier studies, the physical activity questionnaire data can be considered valid.^15–18^ Assessment of the MET index was based on a series of structured questions^15,19^ on leisure-time physical activity (monthly frequency, mean duration, and mean intensity of sessions) and physical activity during commuting. The index was calculated by assigning a MET score to each activity and by calculating the product of that activity: intensity × duration × frequency.^15^ The MET index was expressed as the sum-score of leisure-time physical activity MET-hours per day. To estimate the mean volume of physical activity during the three baseline survey years, the average of the MET index values obtained in 1975, 1981, and 1990 was computed. This new leisure-time mean MET value (LT-mMET index) was then divided into three activity tertiles labelled low (LT-mMET index 0-1.54 MET h/day), medium (1.54-2.92 MET h/day), and high (2.92-26.13 MET h/day) using the same tertiles as in an earlier study.^18^ Twin pairs were classified as discordant for physical activity if one co-twin was in the low-activity tertile and the other co-twin was in the high-activity tertile.

*As other predictors and covariates*, body mass index (BMI), self-reported work-related physical activity, smoking status, use of alcohol and health status in 1990, and socioeconomic status were used. After preliminary analyses, to maximize statistical power for pairwise twin and multivariate analyses, covariates were dichotomized by merging classes not differing for baseline and follow-up physical activity levels.

BMI was calculated based on self-reported height and weight. Work-related physical activity was a categorical variable evaluated with a four-point ordinal scale.^20^ A response to the first option “mainly sedentary work, which requires very little physical activity” was classified as sedentary work, while all other responses (“work that involves standing and walking, but no other physical activity” and more strenuous) were classified as non-sedentary work. Three socioeconomic status categories (white collar, intermediate, and blue collar) were defined by years of education and amount of physical activity at work.^14^ The blue collar and intermediate groups were combined in the analyses because their baseline and follow-up physical activity was similar. Smoking status, originally coded into four categories,^21^ was dichotomized (current daily vs. others) for the main analyses. Alcohol use was expressed as a dichotomous variable of heavy drinking occasions (i.e., consumption of at least six drinks on one occasion) at least monthly.^22,23^ Somatic health status (healthy/not) was defined as having/not having a disease diagnosed by a physician, serious injury/illness, or permanent work disability, according to self-report items in 1990.^14^

### Accelerometer Data Collection and Analysis

Physical activity was measured with a waist-worn, light triaxial accelerometer (Hookie AM20, Traxmeet Ltd, Espoo), which was employed in a previous large population-based study of Finnish adults.^24^ The device and instructions for use were mailed to the participants, who were asked to use the accelerometer during waking hours for 7 consecutive days. Participants mailed the device back to UKK Institute for data analysis, and they were later provided with their own results. The analysis of raw acceleration data was based on novel algorithms that employ the mean amplitude deviation (MAD) of the resultant acceleration during a 6 s epoch and the angle for posture estimation (APE) of the body, metrics that provide a consistent assessment of the intensity of physical activity and separate accurately sedentary and stationary behaviors from any physical activity^25,26^

MAD was also validated through directly measured incident VO_2_ during walking or running on an indoor track.^26^ This strong association allowed for transformation of MAD values to incident energy consumption (MET). The MET values for each minute were calculated as the one-minute exponential moving average of MAD values. According to standard use,^27^ cut-off points for different activities were set as 1.5-3 MET for light activities, 3-6 MET for moderate activities, and over 6 MET for vigorous activities, and corresponding mean daily total times were determined. Mean daily sedentary time was defined as MET under 1.5 during lying down or sitting. Mean daily standing time was analyzed separately. Average daily step count and the most intensive 10-minute period (Peak-10min MET) during the monitoring week were also documented.

Altogether, 791 twin individuals wore the accelerometer for at least 10 hours per day for 4 days. On average, they wore the device 6.73 days (95% confidence interval [CI] 6.69-6.77) and 14:01:44 h:min:sec/day (95% CI 13:56:31-14:04:37). A total of 616 had complete data for calculating MET indices from all of the 1975, 1981, and 1990 questionnaires. No significant differences in MVPA (40.2 min vs. 37.7 min, *P*=.30) and daily steps (6440 vs. 6120, *P*=.23) were seen between these 616 individuals and the 175 individuals who did not have baseline LT-mMET but participated in the accelerometer study.

### Statistical Methods

Descriptive statistics were calculated with bootstrapping (1000 samples unless otherwise noted) and are given as medians and interquartile ranges (IQRs) or 95% confidence intervals (CIs). We used linear regression analyses to define R squared (R^2^) as a measure of variance accounted for. The analyses were done with twins treated as individuals; however, because the observations obtained from twin pairs may be correlated, robust estimators of variance (the cluster option in Stata) were used.^28^ All basic analyses yielding R^2^ values were adjusted for age and sex. To obtain R^2^ only for the studied variable, the variable was entered the model after the basic model and then the difference in R^2^(ΔR^2^) was calculated. Multivariate models were adjusted for BMI, smoking, alcohol, work-related physical activity, health status, and socioeconomic status. Square root-transformation for MVPA, logarithm-transformation for Peak-10min MET, and cubic root transformation for LT-mMET were used for regression analyses because these variables were not normally distributed.

Pairwise analyses among twin pairs (all pairs, DZ pairs, and MZ pairs separately) were done using Wilcoxon matched-pairs signed-rank test for whether pairs discordant for specific baseline characteristics or health habits differed in the objectively measured physical activity variables at follow-up.

Quantitative trait modeling was done using the MET variables from 1975, 1981, and 1990 to analyze whether they were direct risk factors or whether the association with the follow-up physical activity variables was mediated by genetic or other environmental factors. The quantitative trait modeling is described in **eMethods** and **eResults** in the supplementary file, and only the main results are given below.

## Results

### Participant Characteristics and Selection

Mean age of the participants was 48.3 years (range 45.9-51.4) at time of response to the 1990 questionnaire and 72.9 years (range 71.1-75.0) for objective physical activity monitoring. Among those who responded to the baseline LT-mMET questions (1 646 individuals in this age group), the LT-mMET index was similar in those who participated in the follow-up accelerometry study (n=616) and those who did not participate for various reasons (n=1030) (LT-mMET index in MET-h/day 2.65 ± 2.0 vs. 2.69 ± 2.6; men 2.97 ± 2.4 vs. 2.98 ± 3.1; women 2.38 ± 1.6 vs. 2.45 ± 2.0). Baseline participant characteristics by LT-mMET index tertiles are shown in Table 1. Among women, lower LT-mMET index was associated with reduced health, while among men, white collar work was more common in the highest LT-mMET index tertile.

**Table 1.** Baseline Participant Characteristics in 1990 by LT-mMET (1975, 1981, 1990) Tertiles

|  |  | LT-mMET tertile<sup>a</sup> |  |  | *P* value<sup>b</sup> |
| --- | --- | --- | --- | --- | --- |
|  |  | Low | Moderate | High |  |
| All, No. |  | 197 | 221 | 198 |  |
| Men, No. |  | 91 | 93 | 106 |  |
| Women, No. |  | 106 | 128 | 92 |  |
| LT-mMET, median (IQR), MET-h/day |  |  |  |  |  |
| All |  | 0.97 (0.56) | 2.12 (0.76) | 4.11 (2.24) |  |
| Men |  | 0.98 (0.54) | 2.14 (0.71) | 4.52 (3.35) |  |
| Women |  | 0.95 (0.57) | 2.10 (0.76) | 3.79 (1.75) |  |
| Body mass index, median (IQR), kg/m<sup>2</sup> |  |  |  |  |  |
| All |  | 24.8 (3.99) | 24.7 (4.01) | 23.7 (3.28) | .005<sup>c</sup> |
| Men |  | 25.2 (3.33) | 25.5 (3.10) | 24.3 (3.26) | .034 |
| Women |  | 24.2 (4.35) | 23.6 (4.15) | 23.1 (3.46) | .060 |
| Work-related loading, No. (%) |  |  |  |  |  |
| All | Sedentary | 87 (44.6) | 96 (44.0) | 95 (48.7) | .595 |
|  | Non-sedentary | 108 (55.4) | 122 (56.0) | 100 (51.3) |  |
| Men | Sedentary | 40 (44.0) | 45 (48.9) | 52 (49.5) | .722 |
|  | Non-sedentary | 51 (56.0) | 47 (51.1) | 53 (50.5) |  |
| Women | Sedentary | 47 (45.2) | 51 (40.5) | 43 (47.8) | .529 |
|  | Non-sedentary | 57 (54.8) | 75 (59.5) | 47 (52.2) |  |
| Socioeconomic status, No. (%) |  |  |  |  |  |
| All | White collar | 24 (12.5) | 31 (14.4) | 45 (22.8) | .012 |
|  | Others | 168 (87.5) | 185 (85.6) | 152 (77.2) |  |
| Men | White collar | 10 (11.2) | 12 (13.2) | 26 (24.8) | .019 |
|  | Others | 79 (88.8) | 79 (86.8) | 79 (75.2) |  |
| Women | White collar | 14 (13.6) | 19 (15.2) | 19 (20.7) | .368 |
|  | Others | 89 (86.4) | 106 (84.8) | 73 (79.3) |  |
| Cigarette smoking, No. (%) |  |  |  |  |  |
| All | No current smoking | 160 (81.6) | 184 (83.6) | 169 (86.2) | .479 |
|  | Current smoker | 36 (18.4) | 36 (16.4) | 27 (13.8) |  |
| Men | No current smoking | 76 (84.4) | 77 (83.7) | 87 (82.9) | .953 |
|  | Current | 14 (15.6) | 15 (16.3) | 18 (17.1) |  |
| Women | No current smoking | 84 (79.2) | 107 (83.6) | 82 (90.1) | .137 |
|  | Current | 22 (20.8) | 21 (16.4) | 9 (9.9) |  |
| Heavy (high-density drinking occasions) alcohol users, No. (%) |  |  |  |  |  |
| All | No | 151 (77.0) | 177 (81.2) | 153 (77.3) | .500 |
|  | Yes | 45 (23.0) | 41 (18.8) | 45 (22.7) |  |
| Men | No | 59 (65.6) | 59 (64.8) | 69 (65.1) | .995 |
|  | Yes | 31 (34.4) | 32 (35.2) | 37 (34.9) |  |
| Women | No | 92 (86.8) | 118 (92.9) | 84 (91.3) | .259 |
|  | Yes | 14 (13.2) | 9 (7.1) | 8 (8.7) |  |
| Health status, No. (%) |  |  |  |  |  |
| All | Sick | 123 (64.1) | 137 (63.4) | 100 (50.8) | .010 |
|  | Healthy | 69 (35.9) | 79 (36.6) | 97 (49.2) |  |
| Men | Sick | 54 (60.7) | 48 (52.7) | 50 (47.6) | .205 |
|  | Healthy | 35 (39.3) | 43 (47.3) | 55 (52.4) |  |
| Women | Sick | 69 (67.0) | 89 (71.2) | 50 (54.3) | .029 |
|  | Healthy | 34 (33.0) | 36 (28.8) | 42 (45.7) |  |

<sup>a</sup>All descriptive analyses with bootstrapping (1000 samples unless otherwise noted)<sup>b</sup>Rao & Scott Chi-Square test<sup>c</sup>Linear regression cluster for family, with LT-mMET and BMI both used as continuous variables

### Predictors of Later Life Objectively Measured Physical Activity and Sedentary Behavior: Individual-Based Analyses

High baseline LT-mMET index predicted less sedentary behavior (additional R^2^ 2.0% after age- and sex adjustment, *P*=.002), more MVPA (R^2^ 6.9, *P*<.001), more steps (R^2^ 5.6%, *P*<.001) and also higher intensity Peak-10 min MET during the monitoring week (R^2^ 7.5%, *P*<.001) (Table 2, with results also by sex). The LT-mMET index was a stronger predictor of follow-up MVPA than any of the MET values from individual baseline time-points.

**Table 2.** Follow-up Objective Physical Activity Measurements by Mean Baseline LT-mMET index (1975, 1981, 1990) Tertiles

| Activity /<br>inactivity<br>variable<sup>a</sup> | LT-mMET index tertile<sup>b</sup> |  |  | R<sup>2</sup> (%)<sup>c</sup> | P value<sup>d</sup> |
| --- | --- | --- | --- | --- | --- |
|  | Low | Moderate | High |  |  |
| Mean sedentary time/day, median (95% CI), h:min:sec |  |  |  |  |  |
| All | 9:10:22<br>(9:00:59 to 9:18:18) | 8:38:03<br>(8:26:27 to 8:52:01) | 8:38:58<br>(8:20:09 to 9:00:49) | 2.0 | .002 |
| Men | 9:11:19<br>(9:03:35 to 9:43:39) | 8:52:01<br>(8:38:03 to 9:12:51) | 8:43:49<br>(8:23:46 to 9:03:39) | 3.4 | .012 |
| Women | 9:05:05<br>(8:45:39 to 9:22:43) | 8:25:34<br>(8:07:18 to 8:46:19) | 8:23:52<br>(7:58:15 to 9:04:13) | 1.1 | .041 |
| Mean standing time/day, median (95% CI), h:min:sec |  |  |  |  |  |
| All | 1:19:09<br>(1:07:45 to 1:27:03) | 1:28:05<br>(1:21:59 to 1:33:49) | 1:22:10<br>(1:16:32 to 1:29:57) | 0.4 | .110 |
| Men | 1:17:23<br>(1:01:59 to 1:25:40) | 1:21:43<br>(1:12:49 to 1:32:57) | 1:18:18<br>(1:11:52 to 1:29:00) | 1.0 | .090 |
| Women | 1:21:46<br>(1:07:22 to 1:37:36) | 1:30:07<br>(1:24:27 to 1:37:34) | 1:25:45<br>(1:17:15 to 1:35:49) | 0.1 | .509 |
| Mean time of light physical activity/day, median (95% CI), h:min:sec |  |  |  |  |  |
| All | 2:43:53<br>(2:35:57 to 2:54:55) | 2:55:05<br>(2:37:26 to 3:01:21) | 3:01:18<br>(2:46:53 to 3:13:56) | 0.7 | .040 |
| Men | 2:50:12<br>(2:25:46 to 3:01:44) | 2:57:13<br>(2:38:06 to 3:09:59) | 2:52:12<br>(2:31:10 to 3:11:30) | 0.1 | .696 |
| Women | 2:41:43<br>(2:34:47 to 2:55:56) | 2:47:05<br>(2:33:10 to 3:01:21) | 3:09:24<br>(2:55:19 to 3:27:10) | 2.3 | .011 |
| Mean time of moderate-to-vigorous physical activity/day, median (95% CI), h:min:sec |  |  |  |  |  |
| All | 0:26:43<br>(0:22:06 to 0:29:52) | 0:36:31<br>(0:32:16 to 0:40:35) | 0:42:43<br>(0:36:37 to 0:50:49) | 6.9 | <.001 |
| Men | 0:31:31<br>(0:24:52 to 0:38:21) | 0:40:43<br>(0:32:01 to 0:45:47) | 0:47:00<br>(0:34:25 to 0:56:57) | 7.5 | <.001 |
| Women | 0:22:05<br>(0:18:05 to 0:26:57) | 0:34:40<br>(0:28:27 to 0:38:48) | 0:41:00<br>(0:34:18 to 0:49:39) | 6.5 | <.001 |
| Mean daily step count, median (95% CI), No. |  |  |  |  |  |
| All | 5099<br>(4744 to 5706) | 6114<br>(5610 to 6656) | 7072<br>(6245 to 7788) | 5.6 | <.001 |
| Men | 5838<br>(5126 to 6531) | 6519<br>(5958 to 6788) | 7272<br>(6132 to 8340) | 6.7 | <.001 |
| Women | 4753<br>(3964 to 5099) | 5612<br>(5231 to 6656) | 6800<br>(6236 to 7844) | 4.8 | <.001 |
| Peak-10min MET, median (95% CI), MET |  |  |  |  |  |
| All | 3.26<br>(3.17 to 3.38) | 3.57<br>(3.39 to 3.68) | 3.81<br>(3.55 to 3.96) | 7.5 | <.001 |
| Men | 3.35<br>(3.17 to 3.45) | 3.67<br>(3.39 to 3.83) | 3.81<br>(3.43 to 4.18) | 9.9 | <.001 |
| Women | 3.20<br>(2.97 to 3.33) | 3.54<br>(3.30 to 3.66) | 3.81<br>(3.51 to 3.99) | 5.0 | <.001 |

<sup>a</sup>Activity variables calculated based on 1 minute segments<sup>b</sup>Descriptive analyses with bootstrapping (1000 samples)<sup>c</sup>R<sup>2</sup> for LT-mMET index calculated as a difference ( $\Delta R^2$ ) from age and sex model compared to model with LT-mMET + age and sex, indicating the true R<sup>2</sup> of LT-mMET<sup>d</sup>P value calculated with continuous LT-mMET variable from linear regression adjusted for sex and age and cluster for family

Table 3 shows the association between other baseline predictors from 1990 and MVPA at follow-up. High BMI had the strongest association with an additional R^2^ of 10.7% (for details on analyses of daily steps see **eTable 1**).

**Table 3.**
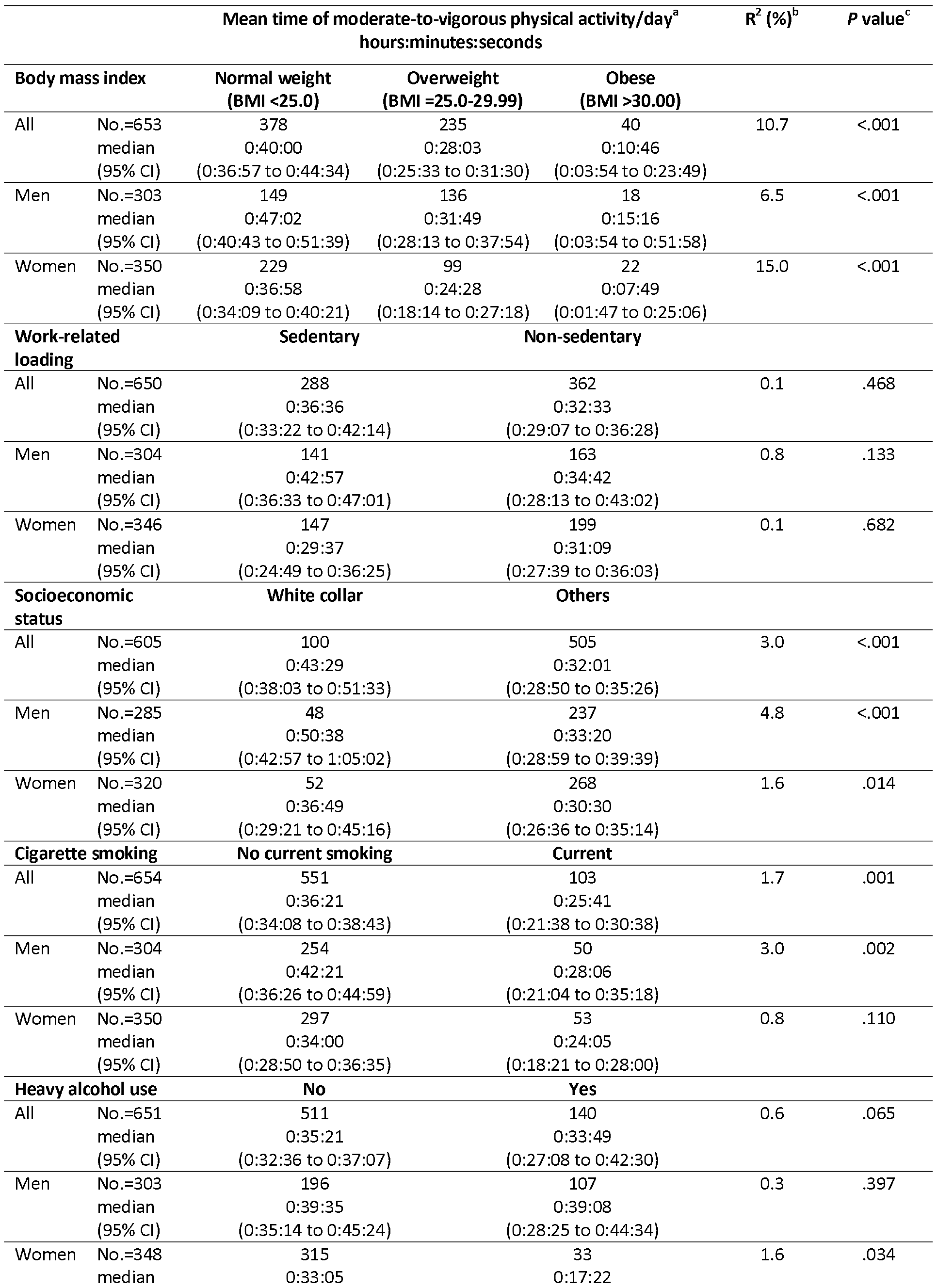
Moderate-to-Vigorous Physical Activity by 1990 Baseline Covariates

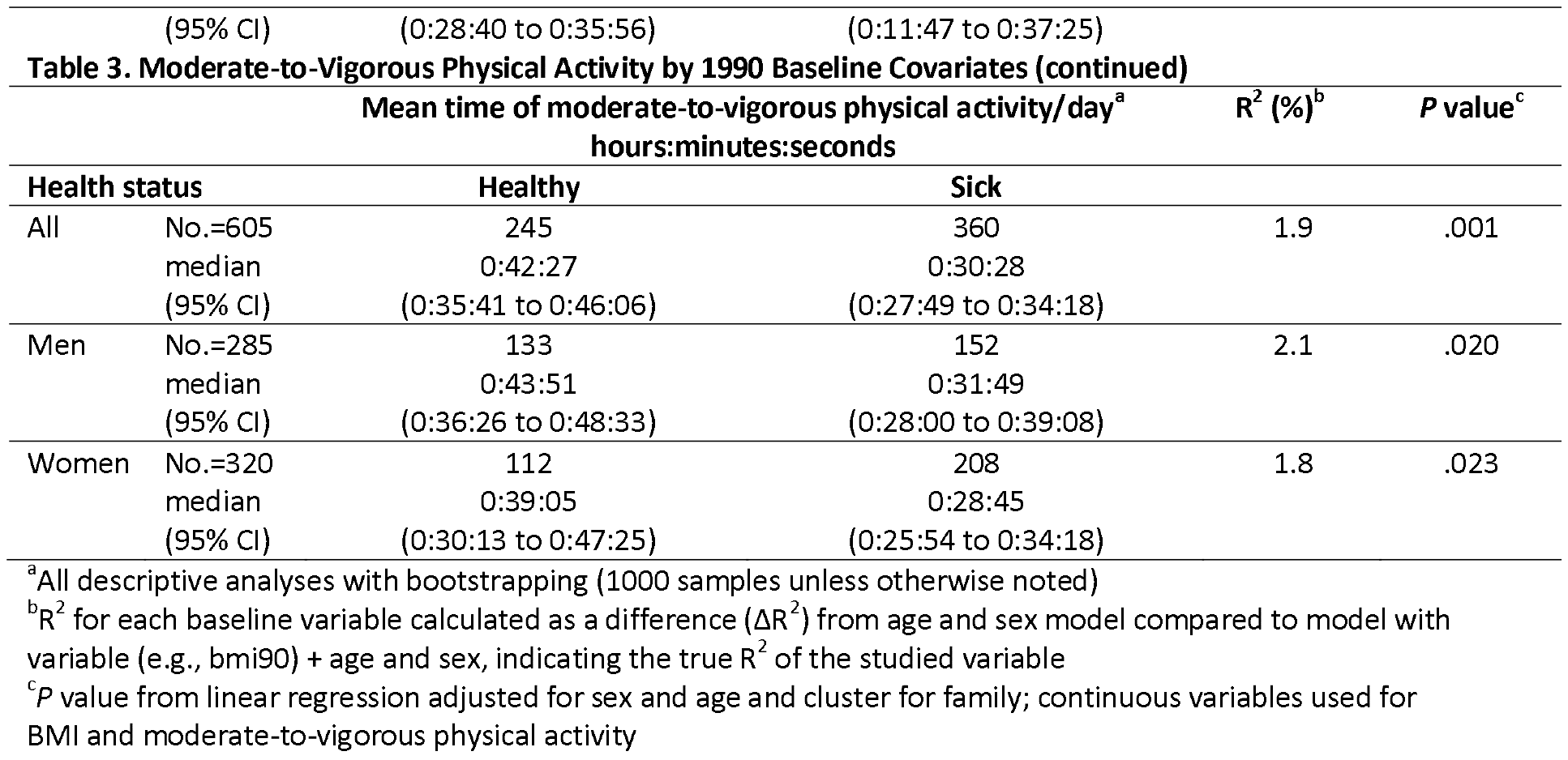

In the multivariate MVPA prediction regression model, with the addition of BMI after age, sex, and LT-mMET index, the R^2^ value increased from 8.4% to 17.2%, and up to 20.3% with smoking, socioeconomic status, and health status also in the model **(eTable 2)**. Use of alcohol and work-related physical activity were not significant contributors when added to this model. Similar models for daily step count showed rather similar results with a slightly lower proportion of variance accounted for.

### Predictors of Later-Life Objectively Measured Physical Activity: Pairwise Analyses

Although there were some trends in the same direction in pairwise analyses among twin pairs discordant for different predictors at baseline, only twin pairs who were discordant for smoking (n=40 discordant pairs; median follow-up MVPA volumes of 25 minutes for current smokers at baseline and 35 minutes for others; *P*=.037) or for health status (n=69 discordant pairs, 30 *vs*. 44 minutes, *P*=.014) differed in their follow-up MVPA volumes (Table 4). For smoking, the difference also was seen for MZ pairs, but for health status, it was seen only for DZ pairs. In the smaller number of socioeconomic status-discordant MZ pairs, lower socioeconomic status predicted less MVPA at follow-up. The trends were similar for daily step count **(eTable 3)**.

**Table 4.**
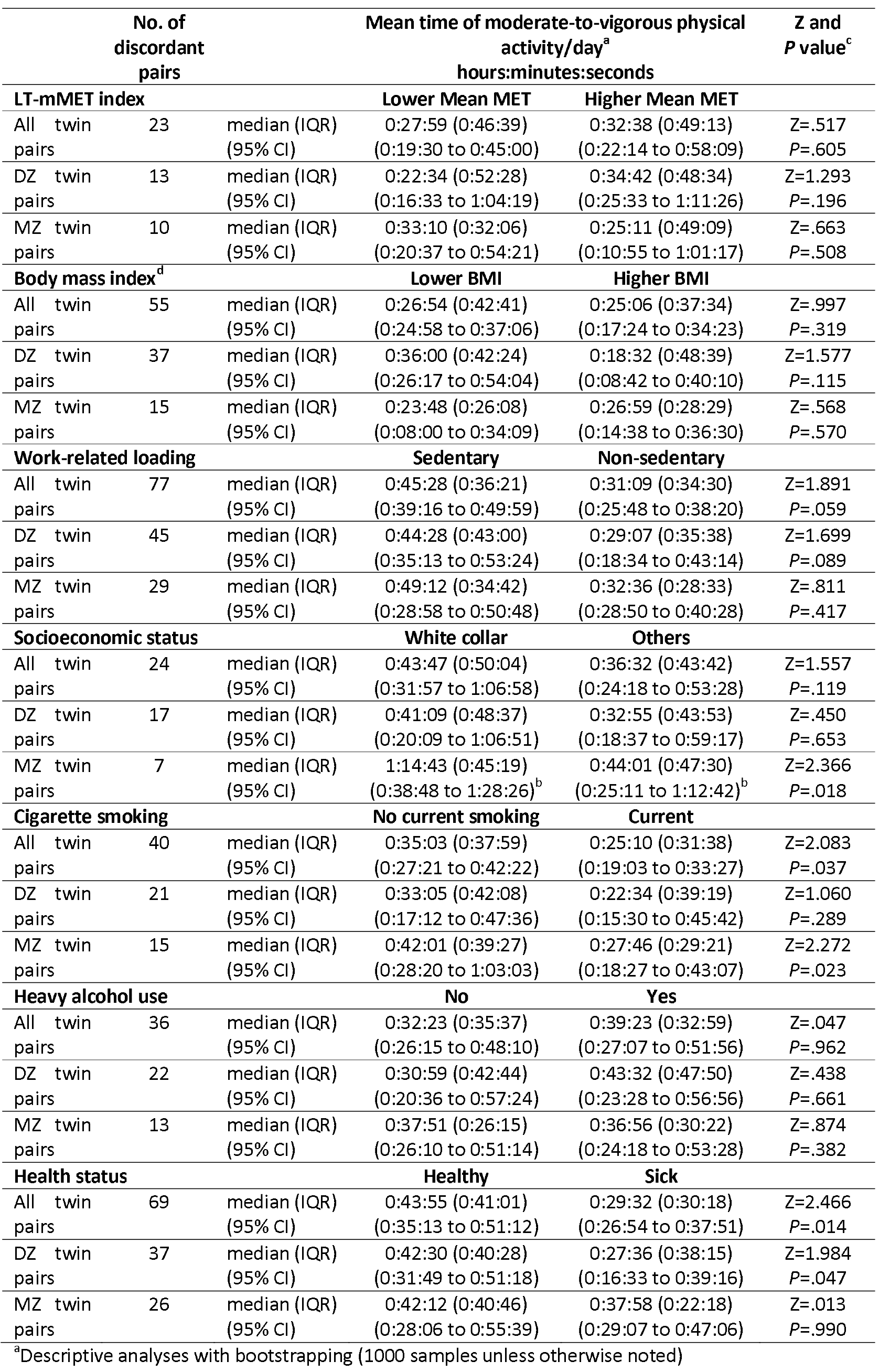
Moderate-to-Vigorous Physical Activity in Twin Pairs Discordant for Different Baseline Characteristics^a^

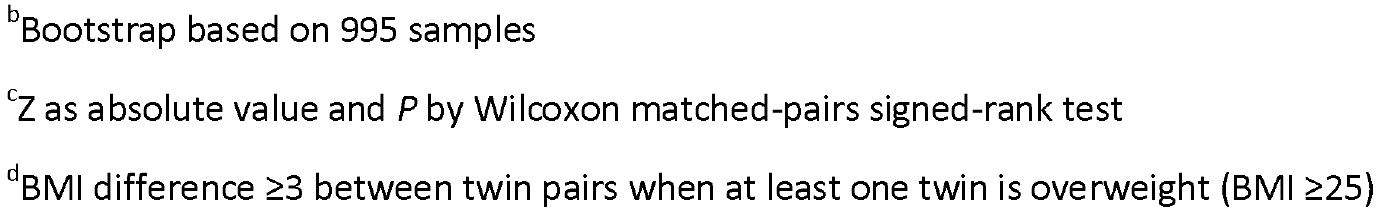

### Mediation Analysis by Quantitative Trait Modeling

Based on quantitative trait models (for more details see Supplementary **eResults, eTables 4-8,** and **eFigure 1**), joint genetic effects mediated the association from baseline MET factor on MVPA and Peak-10min MET. The MET factor was observed to be a direct risk factor for number of daily steps and sedentary behavior (lying and sitting). No relationship was observed of MET factor with standing and light physical activity. In more detail, the broad sense heritability for MVPA was 60% (**eTable 8**). When cross-trait correlation between baseline MET factor and follow-up MVPA was decomposed into genetic and residual parts based on the model where we estimated both the genetic and environmental correlations the estimated cross-trait correlation was 0.35 (95% CI 0.25-0.43) with 82% (53%-100%) contribution from genetic factors.

## Discussion

Younger-age leisure-time physical activity and other covariates explained one fifth (20.3%) of the variation in moderate-to-vigorous activity in older age in this prospective twin cohort study. According to pairwise analyses, much of the association was driven by shared genes underlying mid-life physical activity and later objectively measured activity. Smoking contributed independent of genes.

### Comparison to Other Studies

In line with our findings, high physical activity is associated in cross-sectional or longitudinal designs with high previous physical activity, low BMI, low work-related physical loading, and good health status.^29–31^ In cross-sectional and shorter-term follow-up studies, low physical activity is associated with lower fitness, more frailty, higher disability, and poor health.^32–35^ No long-term randomized trials have addressed whether changes in health behavior in middle age lead to late-life differences in physical activity. Also, observational follow-ups on this topic are rare, and we are not aware of other studies relating long-term leisure-time physical activity differences in younger adulthood to objectively measured physical activity/inactivity in later years.^36^

In our individual-based analyses, we found significant predictors for later-life physical activity, but could not replicate all of the results in pairwise analyses among the predictor-discordant MZ twin pairs. The outcome is a reminder that genetic or other familial factors may explain why associations are often seen between younger-age physical activity and later-age health-related factors and, consequently, mobility.

Smoking at baseline also predicted less MVPA at follow-up in pairwise analysis among MZ twin pairs, which is evidence for an association not explained by genetic factors. These results are in line with our earlier finding that MZ twin pairs discordant for smoking show a clear difference in overall mortality while pairs discordant for physical activity participation do not.^20,37^ Our quantitative trait modelling was in agreement with the results of the pairwise analyses. Smoking affects both pulmonary and cardiovascular health and increases systemic inflammation, all of which may decrease the ability to exercise. We cannot exclude the possibility that smoking is also a marker for other lifestyle factors that predict less physical activity.

The strengths of our study include that we had physical activity data from three different baseline time-points, a nationally representative large twin cohort, very long-term prospective data, and novel valid analysis of the follow-up physical activity and sedentary behavior profile.^27^

### Limitations

Our study has several limitations. Our baseline predictor assessments relied on self-reported questionnaire data. We lack comprehensive data on dietary factors or clinical examinations at baseline. Although our study was large enough for the individual-based analyses, the number of MZ twin pairs discordant for some of the predictors was quite low providing only moderate statistical power for some analyses. At follow-up, most twins were community dwelling, so individuals with severe mobility limitations were rare.

### Conclusions

Our follow-up study among twins showed that middle-age low leisure-time physical activity, obesity, smoking, low socioeconomic status, and health problems predicted low MVPA at older age in individual-based analyses. According to pairwise analyses, smoking seemed to causally predict less physical activity in later years while other associations were more likely attributable to shared genetic factors and childhood environment.

## ARTICLE INFORMATION

**Author Affiliations: See page 1**

### Author Contributions

Dr Kujala, as principal investigator, had full access to all data in the study and takes responsibility for the integrity of the data and the accuracy of the data analysis.

*Study concept and design:* Waller, Kaprio, Sievänen, Kujala.

*Acquisition, analysis, or interpretation of data:* All authors.

*Drafting of the manuscript:* Waller, Kujala.

*Critical revision of the manuscript for important intellectual content:* All authors.

*Statistical analysis:* Waller, Vähä-Ypyä, Törmäkangas (quantitative trait modeling), Hautasaari, Kujala.

*Administrative, technicalor material support:* Rinne, Kaprio, Sievänen, Kujala.

*Study supervision:* Rinne, Kaprio, Sievänen, Kujala.

